# Neural mechanisms of context-dependent segmentation tested on large-scale recording data

**DOI:** 10.1101/2021.04.25.441363

**Authors:** Toshitake Asabuki, Tomoki Fukai

**Affiliations:** Okinawa Institute of Science and Technology, Onna-son, Okinawa 9040495, Japan

## Abstract

The brain performs various cognitive functions by learning the spatiotemporal salient features of the environment. This learning likely requires unsupervised segmentation of hierarchically organized spike sequences, but the underlying neural mechanism is only poorly understood. Here, we show that a recurrent gated network of neurons with dendrites can context-dependently solve difficult segmentation tasks. Dendrites in this model learn to predict somatic responses in a self-supervising manner while recurrent connections learn a context-dependent gating of dendro-somatic current flows to minimize a prediction error. These connections select particular information suitable for the given context from input features redundantly learned by the dendrites. The model selectively learned salient segments in complex synthetic sequences. Furthermore, the model was also effective for detecting multiple cell assemblies repeating in large-scale calcium imaging data of more than 6,500 cortical neurons. Our results suggest that recurrent gating and dendrites are crucial for cortical learning of context-dependent segmentation tasks.

## Introduction

The ability of the brain to learn hierarchically organized sequences is fundamental to various cognitive functions such as language acquisition, motor skill learning, and memory processing [1-8]. To adequately process the cognitive implications of sequences, the brain has to generate context-dependent representations of sequence information. For instance, in language processing the brain may recognize “nueron” as a misspelling of “neuron” if the brain knows the word “neuron” but not the word “nueron”. However, the brain recognizes “affect” and “effect” as different words even if the two words are very similar. “Break” and “brake” are also different words although these words combine the same letters in different serial orders. The brain can also recognize the same word presented in different temporal lengths. All these examples suggest the inherent flexibility of context-dependent sequence learning in the brain. However, the neural mechanisms underlying this flexible learning, which occurs in an unsupervised manner, remain elusive.

Segmentation or chunking of sensory and motor information is at the core of the context-dependent analysis of hierarchically organized sequences [9-12]. However, little is known about the neural representations and learning mechanisms of hierarchical sequences. Recently, neurons encoding long-range temporal correlations in the song structure were found in the higher vocal center of songbirds [13]. These neurons responded differently to the same syllables (the basic elements of bird song) depending on the preceding phrases (constituted by several syllables) or the succeeding phrases. Different responses of the same neurons in different sequential contexts were also found in the monkey supplementary motor area [14,15]. Hierarchical sequences are often assumed to mirror the hierarchical organization of brain regions. However, the human premotor cortex jointly represents movement chunks and their sequences [16] and linguistic processing in humans also lacks an orderly anatomical representation of sequential context [17].

Unsupervised, context-dependent segmentation is difficult in computational models. Recurrent network models can generate rich sequential dynamics, but these networks are typically trained by a supervised method. Spike-timing-dependent plasticity was used for unsupervised segmentation of input sequences in a recurrent network model [18]. However, while the model worked for simple hierarchical spike sequences, it could not learn context-dependent representations for overlapping spike sequences. Single-cell computation with dendrites could solve a variety of temporal feature analysis including the unsupervised segmentation of hierarchical spike sequences [19], supporting the role of dendrites in sequence processing [20]. However, context-dependent segmentation was also difficult for this model. Although the segmentation problem has been partially solved, recurrent connections alone or dendrites alone are insufficient for solving the difficult segmentation tasks such as exemplified in the beginning of this article.

Here, we demonstrate that a combination of dendritic computation and a recurrent gating dramatically improves the ability of neural networks to context-dependently segment hierarchical spike sequences. Our central hypothesis is that recurrent synaptic input multiplicatively regulates the degree of gating of instantaneous current flows from the dendrites to the soma. As suggested previously [19,21], the dendrites in this model predict the somatic responses. We derive an optimal learning rule for afferent and recurrent synapses to minimize a prediction error. The resultant dendrites generally learn redundant representation of multiple sequence elements while recurrent input learns to selectively gate the dendritic activity suitable for the given context. Recurrent networks with gating synapses were recently used in supervised sequence learning [22].

There is an increasing need for efficient methods to detect and analyze the characteristic spatiotemporal patterns of activity in large-scale neural recording data. We demonstrate that the proposed model can efficiently detect cell-assembly structures in large-scale calcium imaging data. We show two example cases of such analysis in the hippocampus and visual cortex of behaving rodents. In particular, the latter dataset contains the activity of tremendously many neurons (∼ 6,500), and analyzing the fine-scale spatiotemporal structure of activity patterns is computationally costly and difficult for any other methods. In contrast, the data size hardly affected the performance and speed of learning in our model. Surprisingly, the efficiency was even somewhat higher for larger data sizes. These results highlighted the crucial role of recurrent gating in amplifying the weak signature of cell assembly structure detected by the dendrites.

## Results

### The role of recurrent-driven gating in complex segmentation tasks

Our recurrent network model consists of two-compartment neurons with somatic and dendritic compartments (Fig.1a, b). The dendritic components receive hierarchically structured afferent input and recurrent synaptic input, and the sum of these inputs drives the activity of the dendritic component. A similar two-compartment model without recurrent inputs has been studied in segmentation problems [19]. Here, we hypothesize that recurrent synaptic input multiplicatively amplifies or attenuates a current flow from the dendrite to the soma in an input-dependent fashion: The stronger the recurrent input, the larger the dendro-somatic current flow. This “gating” effect is described by a non-linear function of recurrent input to the neuron and controls the instantaneous impact of dendritic activity on the soma (Fig. 1c). We will show below that the recurrent-driven gating plays a pivotal role in the learning of flexible segmentation. Unless otherwise stated, below the results are shown for network models with multiplicative recurrent inputs but no additive ones. In this setting, afferent inputs can evoke large somatic responses if and only if both dendritic activity and gating effect are sufficiently strong. A network model with both additive and multiplicative recurrent inputs will be considered later.

**Figure 1.**
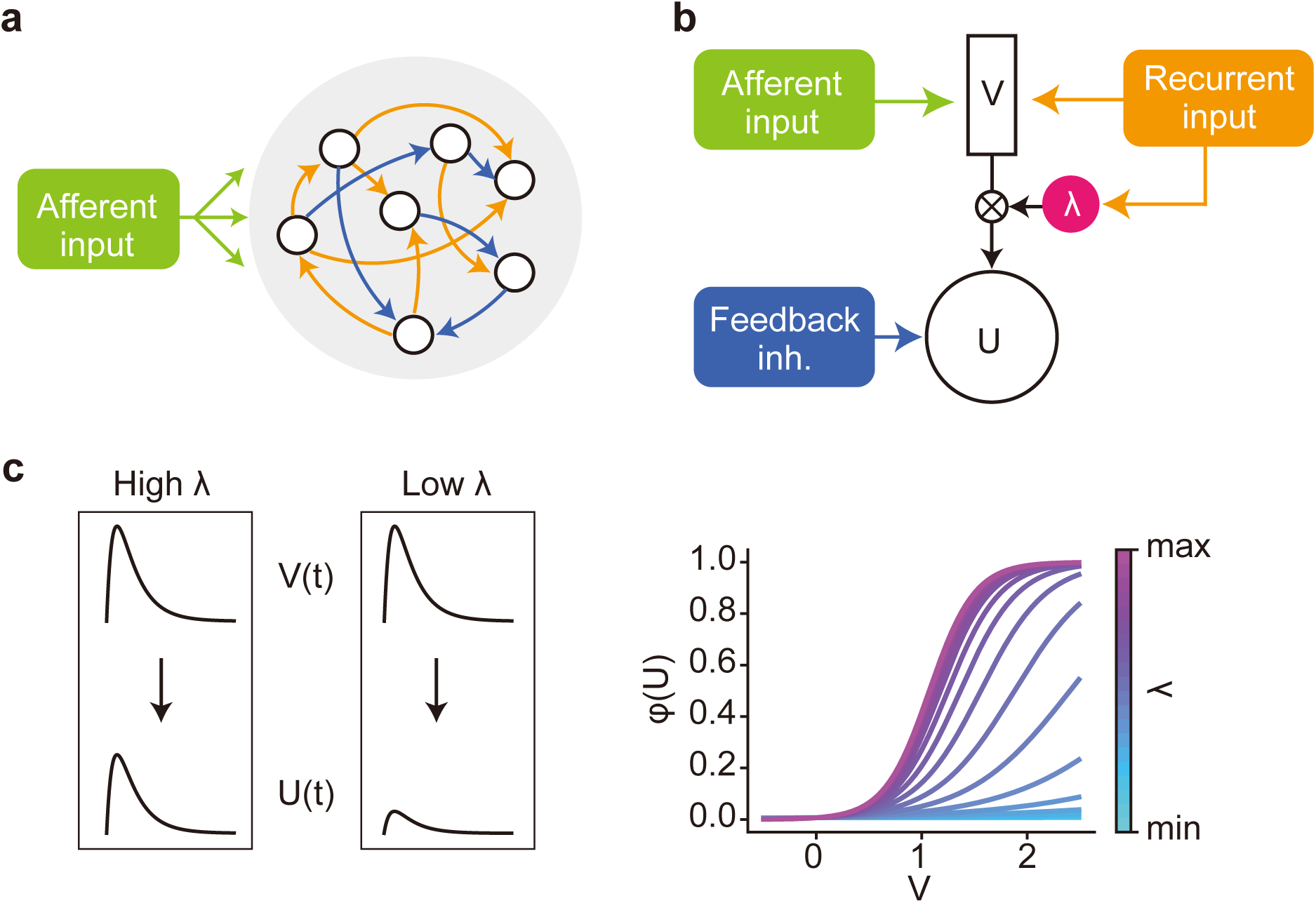
A recurrent gated network of compartmentalized neurons. **a**, A network of randomly connected compartmentalized neurons is considered. **b**, Each neuron model consists of a somatic compartment and a dendritic compartment. Unless otherwise specified, the dendrite only receives afferent inputs. The somatic compartment integrates the dendritic activity and inhibition from other neurons. Gating factor λ is determined by recurrent inputs and regulates the instantaneous fraction of dendritic activity propagated to the soma. **c**, The effect of gating factor on the somatic response is schematically illustrated (left). The relationship between the dendritic potential and somatic firing rate is shown for various values of gating factor (right).

We first demonstrate the segmentation of two spike pattern sequences (chunks) repeated in input spike trains (Fig. 2a). Each chunk was a combination of three fixed spike patterns out of the total five: “A”, “B”, “C”, “D” and “E”, where the component pattern “E” appeared in both chunks. Therefore, the two chunks were mutually overlapped. We fixed the average firing rate of each input neuron at 5 Hz over the entire period of simulations. As training proceeded, the network generated two distinct cell assemblies, each of which selectively responded to one of the chunks (Fig. 2b, c). Notably, each cell assembly responded to the pattern “E” in a preferred chunk of the cell assembly but not in a non-preferred chunk (Fig. 2c, d). A network with a constant gating function trained on the same afferent input failed to discriminate the pattern “E” in different chunks and consequently could not learn the chunks (Supplementary Fig. 1). The result suggests the crucial role of the recurrent-driven gating in the segmentation task.

**Figure 2.**
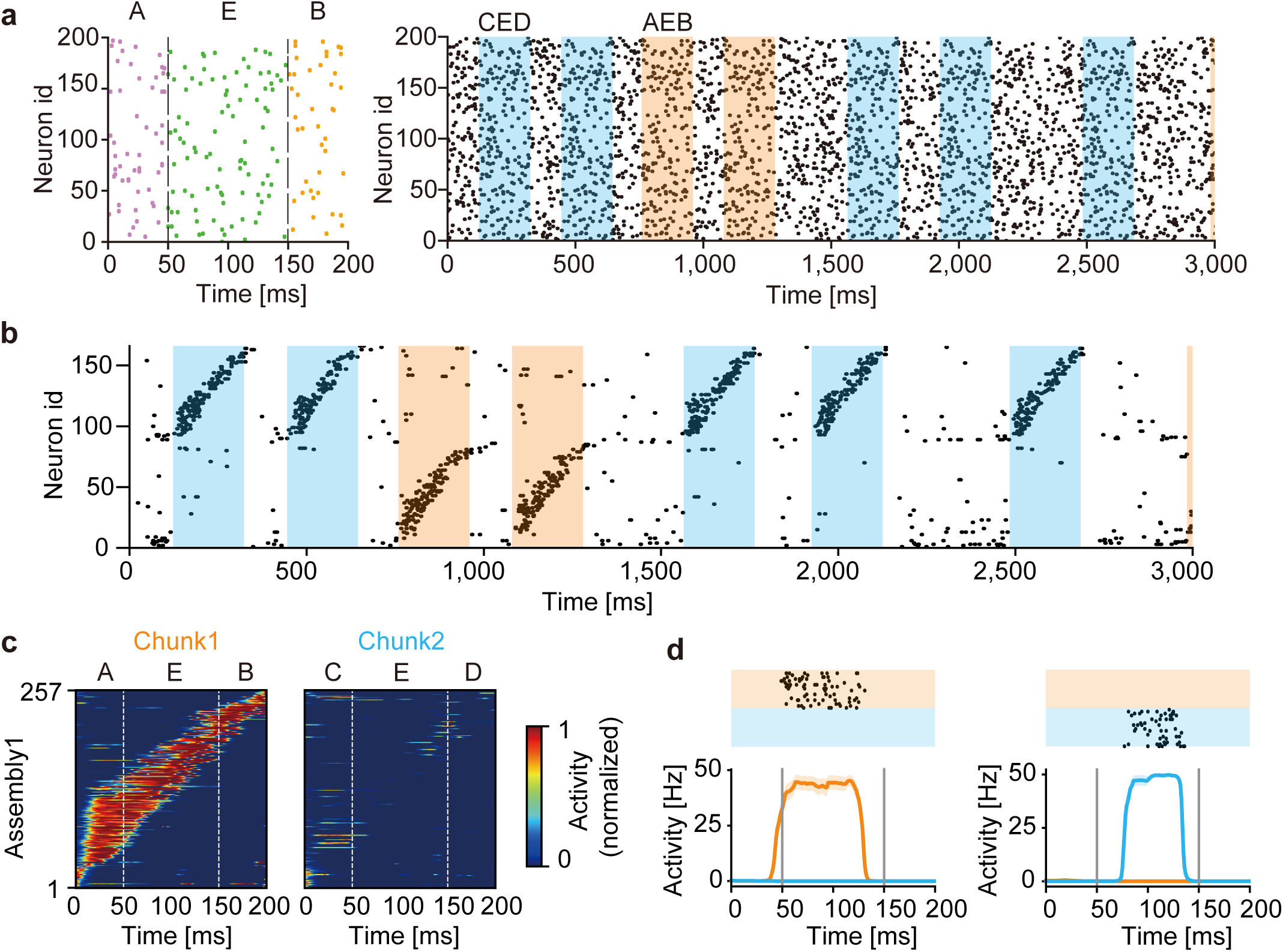
Learning of overlapping chunks. **a**, Two chunks “AEB” (orange shade) and “CED” (blue shade) were repeated in input Poisson spike trains (left). The chunks were separated by random spike trains with variable lengths of 50 to 400 ms (unshaded). All neurons had the same firing rate of 5 Hz. Example spike trains during the initial 3 seconds are shown (right). In each chunk, the component patterns “A”, “B”, “C”, and “D” were 50 ms-long and the shared component “E” was 100 ms-long. **b**, Output spike trains of the trained network model. Neurons were sorted according to their onset response times, and only 160 out of the total 500 neurons are shown for the visualization purpose. A selective cell assembly emerged for each chunk. **c**, The average responses of “AEB”-selective assembly to chunks “AEB” (left) and “CED” (right) are shown. The responses were averaged over 20 presentations of the chunks and normalized by the maximal response to the preferred chunk (i.e., chunk 1). **d**, Reponses of a “AEB”-selective neuron (left) and a “CED”-selective neuron in the trained network are shown.

The above results suggest that this model can discriminate the context of sequences (i.e., the relationship between “E” and other component patterns in a chunk). However, it is also possible that the model separated the overlapping chunks merely relying on the component patterns that were not common between the two chunks and/or on the nonhomogeneous occurrence probabilities among the component patterns. To exclude these possibilities, we examined the case where different chunks shared all component patterns with equal frequencies (Fig. 3a, left). In other words, the same component patterns occurred in different chunks in different orders (i.e., “ABCD”, “DCBA”, “BDAC”). During learning, the model was exposed to irregular spike trains recurring the three chunks intermittently (Fig. 3a, right). The model developed distinct cell assemblies responding selectively to one of these chunks (Fig. 3b, Supplementary Fig. 2a and 2b), thus successfully discriminating the same components belonging to different chunks.

**Figure 3.**
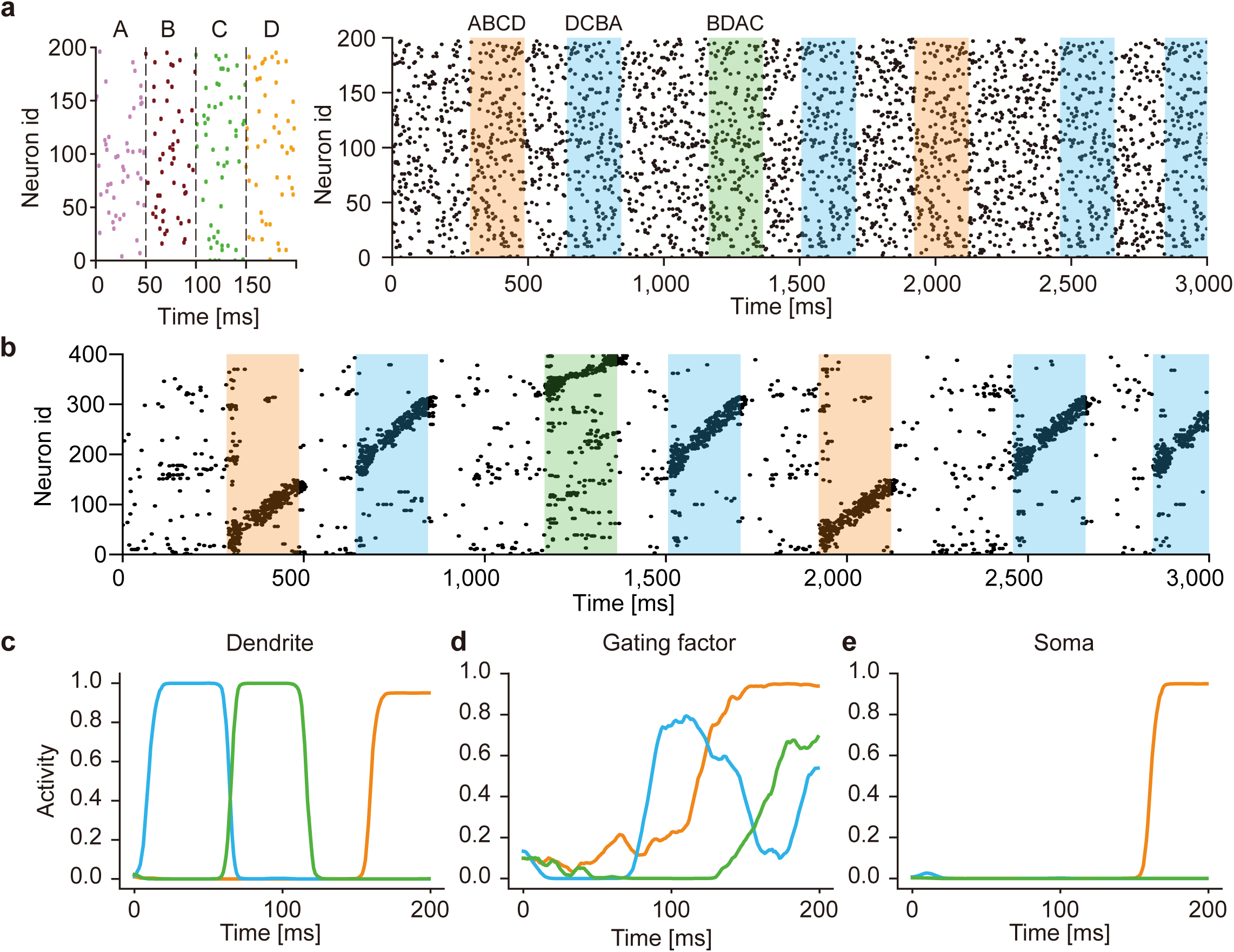
Learning of complex spike sequences with recurrent gating. **a**, Three chunks were presented, which consists of four identical component patterns with its specific order (left). The example input spike train during first three seconds are shown (right). The regions filled in orange, blue and green represent the times when chunks “ABCD”, “DCBA” and “BDAC” were presented, respectively. All neurons generate Poisson spike with the constant firing rate of 5Hz. Random spike sequences were presented in the unshaded areas. **b**, Output spike trains of the trained network are shown. Neurons were sorted according to their onset response times, and only 400 out of 1200 neurons are shown for the visualization purpose. **c, d, e**, The time evolution of dendritic activity, gating factor, and somatic response are shown in a neuron. Orange, blue and green traces represent the responses to its preferred component pattern “D” when the corresponding chunks were presented. The activities were averaged over 20 trials.

### The network mechanism of context-dependent gating

In this model, the recurrent-driven gating provides context-dependent signals necessary for segmenting overlapping chunks. To gain an insight into the role of recurrent gating in learning, we investigated how the somatic and dendritic compartments and gating factors of individual neurons behave during training. The dendritic compartments of these neurons responded to a preferred component pattern irrespective of which chunk the component appeared, showing that the dendrites were unable to discriminate the same component pattern as shared by different chunks (Fig. 3c). In contrast, the gaiting factors responded differently to the same component pattern appearing in different chunks depending on the preceding component patterns (Fig. 3d). This selective gating is thought to arise from the memory effect generated by recurrent synaptic input. As a consequence, the somatic compartment could selectively respond to a particular chunk that strongly activated the gating factor during the presentation of the preferred component pattern (Fig. 3e). Similar results are shown for other neurons in the trained network (Supplementary Fig. 2c)

Our model developed low-dimensional representations that strongly reflect the temporal structures of chunks. The principal component analysis (PCA) of the network responses to the overlapping chunks shown in Fig. 2 revealed that a smaller number of eigenvectors explained a larger cumulative variance as the training progressed (Fig. 4a). At different stages of learning, the low-dimensional trajectories differently represented the chunks. Before learning, the two chunks and unstructured input segments (i.e., random spike trains) occupied almost the same portions of the low-dimensional trajectories (Fig. 4b, left). At the mid stage of learning, the portions of the chunks grew while those of the random segments shrank, but the trajectories evoked by the presentation of the shared pattern “E” were not well separated (Fig. 4b, middle). After sufficient learning, the trajectories were completely separated (Fig. 4b, right). As previously shown, recurrent gating crucially contributed to this separation. Indeed, the network generated almost overlapping trajectories for the pattern “E” if we fastened recurrent gating during learning and test (Fig. 4c) or if we trained the model with recurrent gating but fastened it during test (Fig. 4d). Two trajectories for the overlapping chunks were not clearly separable due to large fluctuations if we randomly shuffled the learned recurrent connections to destroy their connectivity pattern (Fig. 4e). Thus, the context-dependent gating depends crucially on the learned fine structure of recurrent connections.

**Figure 4.**
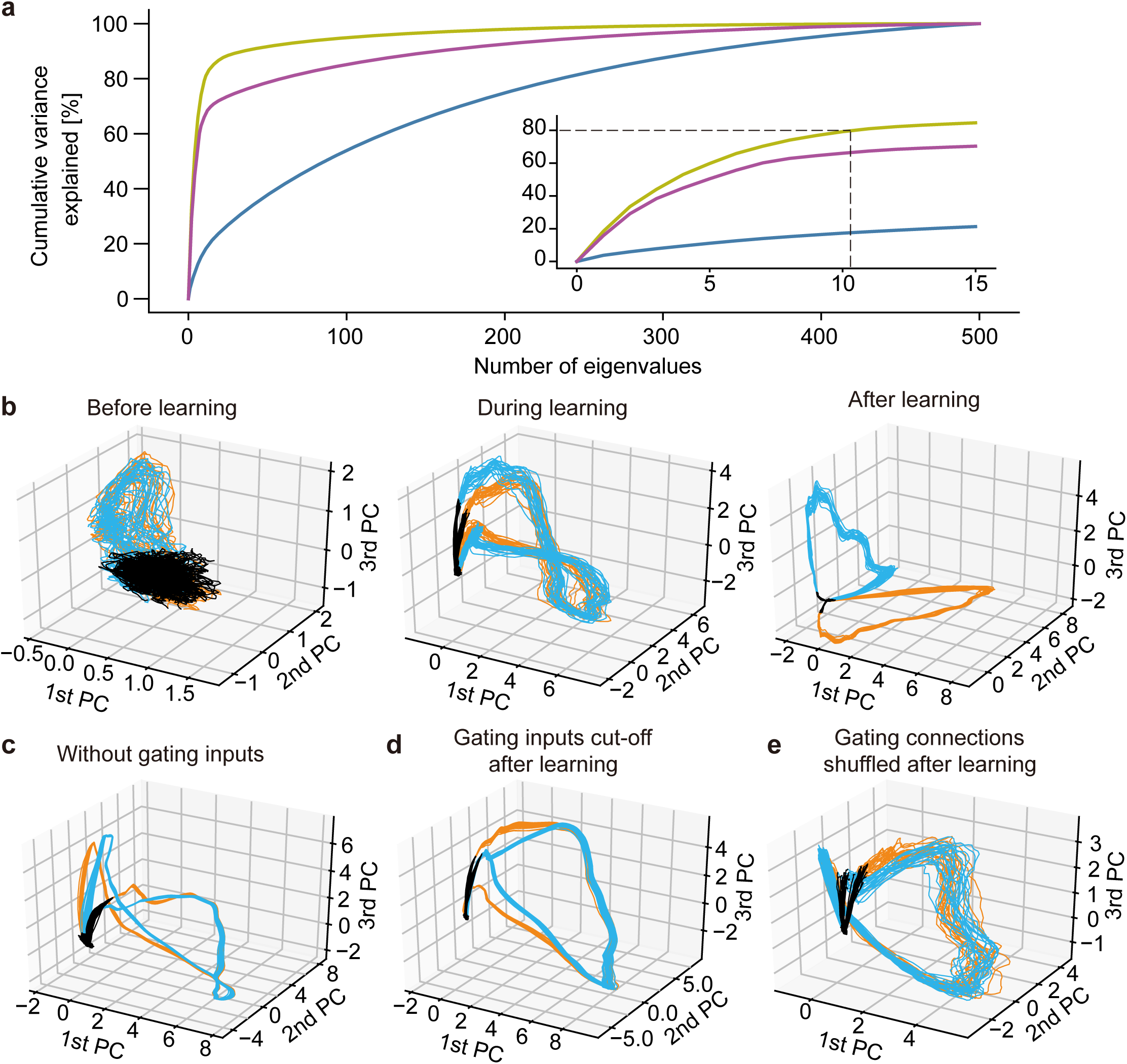
Principal component analysis of the trained network. **a**, Cumulative variance explained of the PCs of the activities of before (blue), during (purple), and after (yellow) learning. Inset is an expanded view for major eigenstates. **b**, The PCA-projected trajectories of network activity before, during, and after training are shown in the space spanned by PC1 to PC3. The network was trained with the same task as in Figure 2. The black, orange, and blue trajectories represent the periods during which random spike input, chunk 1 and chunk 2 were presented, respectively. **c**, Recurrent gating was fixed in all neurons during the whole simulation. **d**, The gating factor was clamped after the network in **b** were trained. **e**, Recurrent connections were randomly shuffled after the network in **b** were trained.

While the network model could learn noisy chunks as far as jitters in spike times were not too large (Supplementary Fig. 3a and 3b), the magnitude of jitters strongly influenced learning speed. This was indicated by the slow saturation of normalized mutual information (see Methods) between network responses and the true labels of chunks during learning (Supplementary Fig. 3c). The normalized mutual information took near the maximum value (≈ 1) as far as the variance of jitters fell within the length of chunks (50 ms). This information dropped rapidly beyond the chunk length (Supplementary Fig. 3d).

Just like the brain can recognize a learned sequence irrespective of the length of its presentation, a learned pattern is detectable for the network even if the pattern is presented with a length different from the learned one (Fig. 5a, b). We quantified the similarity of network responses to otherwise same input patterns with different lengths by calculating the rank-order correlation between responses to stimuli presented with three different lengths. In the network that learned the original pattern, the similarity increased significantly for all three durations of stimulus presentation (Fig. 5c), suggesting that our model learns the manifold of temporal spike patterns rather than individual specific patterns. The robustness shown above raises a question about whether the present model can discriminate precise temporal spike patterns. Indeed, the network model clearly discriminated between similar but different input patterns when the inputs were learned as separate chunks. To study this, we trained the network model with random spike trains involving a repeated temporal pattern and stimulated the learned model with the original pattern (Fig. 5d, top) and its time-reversal version (Fig. 5d, bottom). The cell assembly that learned to detect the original pattern also responded to the reversed pattern in a reversed temporal order, meaning that the different temporal spike patterns were not discriminable in this case (Fig. 5e). Interestingly, the same network model trained with both original pattern and time-reversed pattern self-organized distinct cell assemblies selective for the individual patterns (Fig. 5f). This result may account for discrimination between “break” and “brake” when these words were learned as separate entities.

**Figure 5.**
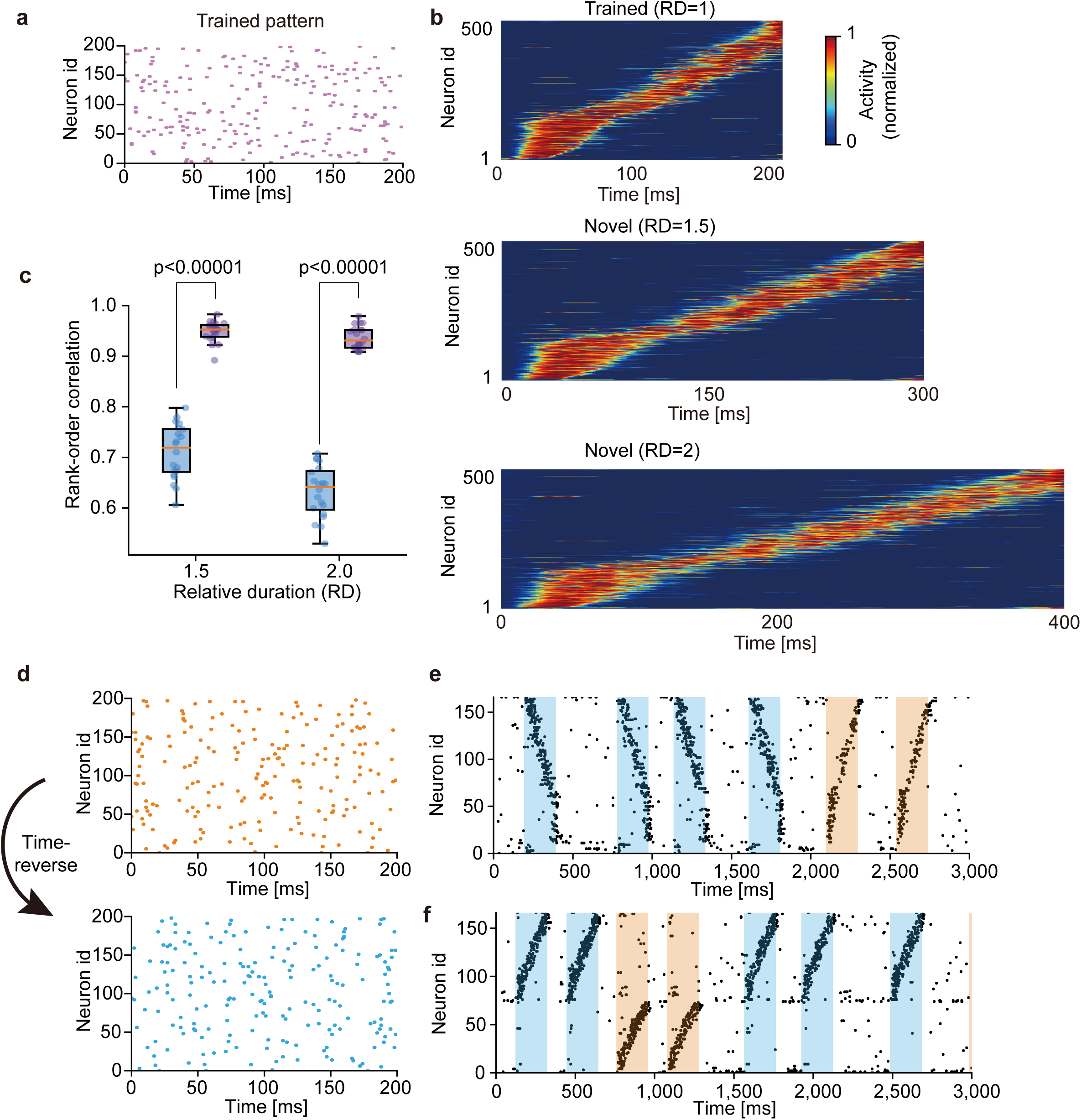
Context-dependent learning of sequence information. **a**, For testing on time-warped patterns, the network was trained on random spike trains embedding a single pattern. **b**, The trained network responded sequentially to the original and stretched patterns with two untrained lengths (i.e., the relative durations RD of 1.5 and 2). **c**, Similarities of sequential order between the responses to the original and two untrained patterns were measured before (blue) and after (purple) learning. Independent simulations were performed 20 times, and p-values were calculated by two-sided Welch’s t-test. **d**, A time-inverted spike pattern (bottom) was generated from a original pattern (top). **e**, The network was exposed to the original pattern in **d** during learning, and its responses were tested after learning for both original and time-inverted patterns. Both patterns activated a single cell assembly. **f**, The network was exposed to both original and time-inverted patterns during learning as well as testing. Two assemblies with different preferred patterns were formed. For the visualization purpose, only 160 out of 500 neurons are shown in **e** and **f**.

### Cell assembly detection in large-scale calcium imaging data

A virtue of our model is its applicability to analyzing large-scale neural recoding data. We show this in two calcium imaging data. The first data contains the activities of 452 hippocampal CA1 neurons recorded from mice running back and forth along a linear track between two rewarded sites (Fig. 6a, top) [23]. Repetitive sequential activations of place cells were reported previously in the data (https://github.com/zivlab/island. For the use of our model, we binarized the data by thresholding activity of each neuron at the 50% of its maximal intensity (Fig. 6a, bottom). After training, model neurons detected groups of input spike trains that tended to arrive in sequences, each of which was preferentially observed at a particular position of the track in a particular direction of run (Fig. 6b). Sorting the activities of hippocampal neurons according to the sequential firing of model neurons (Methods) revealed place-cell sequences without referring to the behavioral data (Fig. 6c).

**Figure 6.**
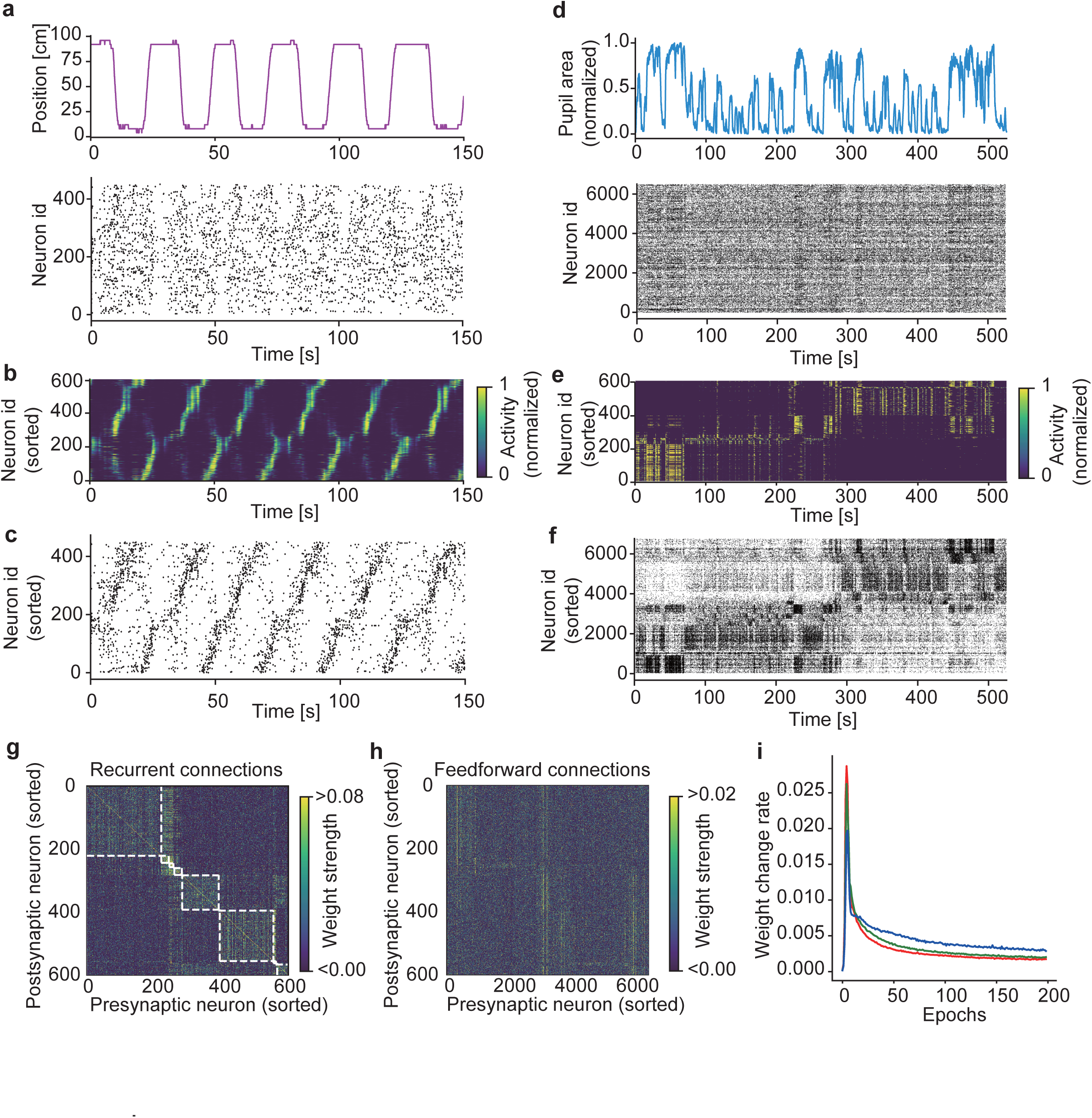
Detecting salient activity patterns in calcium imaging data. **a**, The positions of a mouse (top) on a linear track and calcium imaging data of activity of 452 hippocampal CA1 neurons (bottom) were obtained from previously recorded data [23]. **b**, The learned activities of model neurons were sorted according to their onset response times. **c**, Each CA1 neuron was associated with a model neuron having the highest mutual correlation with the CA1 neuron. Then, the CA1 neurons were sorted according to the serial order of model neurons shown in **b. d**, The time course of normalized pupil area (top) and simultaneously recorded activities of 6,532 visual cortical neurons (bottom) were calculated from previously recorded data [24,25]. **e**, Activity of a trained network model was sorted according to their onset response times (Methods). **f**, Activities of the cortical neurons were sorted as in **c. g**, Sorted recurrent connections are shown. The white dashed lines represent groups of neurons showing co-activation. **h**, Same as in **g**, but sorted feedforward connections are shown. **i**, Learning curves over 200 epochs for various size of input neurons are shown. Red, blue and green traces show learning curves with number of input neurons 1, 2/3, 1/3 times greater than original 6,532 neurons. The weight change rate was calculated as the ratio of the sum of the absolute values of synaptic changes to the sum of the absolute values of all synapses.

Our second example is from the visual cortex in mice running on an air-floating ball [24]. The dataset [25] contains the activity of 6,532 neurons recorded by two-photon calcium imaging from visual cortex as well as behavioral data (facial movements) monitored simultaneously with an infrared camera (Fig. 6d) (https://figshare.com/articles/dataset/Recordings_of_ten_thousand_neurons_in_visual_cortex_during_spontaneous_behaviors/6163622/4. Due to the large data size, detecting cell assemblies is computationally challenging in this dataset. After training, the network model formed several neural ensembles, each of which displayed distinct spatiotemporal response patterns (Fig. 6e). Interestingly, these neural ensembles showed their maximal responses at different periods of time, and the pupil area also changed its maximal size depending on active neural ensembles (Fig. 6d and 6e). By sorting cortical neurons according to the response patterns of model neurons (Methods), we could find the repetition of distinct cell assemblies in the visual cortex (Fig. 6f). The result revealed that active cell assemblies were changed between the early (< 280-290 s) and late epoch of spontaneous behavior. To find the cell-assembly structures, we sorted recurrent connections by grouping co-activated model neurons (Fig. 6g: see Methods). In contrast, such structures were unobvious in the weight matrix of afferent synapses (Fig. 6h). Importantly, without recurrent gating (in this case, the model is equivalent to the previous feedforward network [19]) the cell-assembly structures detected were vague, indicating the crucial contribution of recurrent gating to cell assembly detection (Supplementary Fig. 4). Unexpectedly, the time necessary for learning little changed with data size, or the time was even slightly shorter for larger data sizes (Fig. 6i). Presumably, this unintuitive result was because each cell assembly was represented by more neurons in larger data [19].

### Redundant information representations on dendrites

Evidence from the visual cortex [26], retrosplenial cortex [27], and hippocampus [28] suggested that the representations of sensory and environmental information in cortical neurons are more redundant on the dendrites compared to the soma. The dendrites can have multiple receptive fields while the soma generally represents only one of these receptive fields. The soma is likely to access information represented in a subset of the dendritic branches that share the same receptive field. A similar redundant coding occurs in the present somato-dendritic sequence learning. To show this, we constructed a recurrent network of neurons having three dendritic components and simulated how the model learns orientation tuning. The individual dendritic branches were assumed to undergo independent recurrent gating and mutual competition through softmax (Fig. 7a). During learning, the competition suppressed the dendritic activities that were less correlated with the somatic responses. In the self-organized network, the somatic compartments acquired unique preferred orientations (Fig. 7b). In contrast, dendritic branches displayed different preferred orientations in some neurons (Fig. 7c, d). Thus, the learning rule and recurrent gating proposed in this study possibly underlie the somatic selection process of redundant dendritic representations.

**Figure 7.**
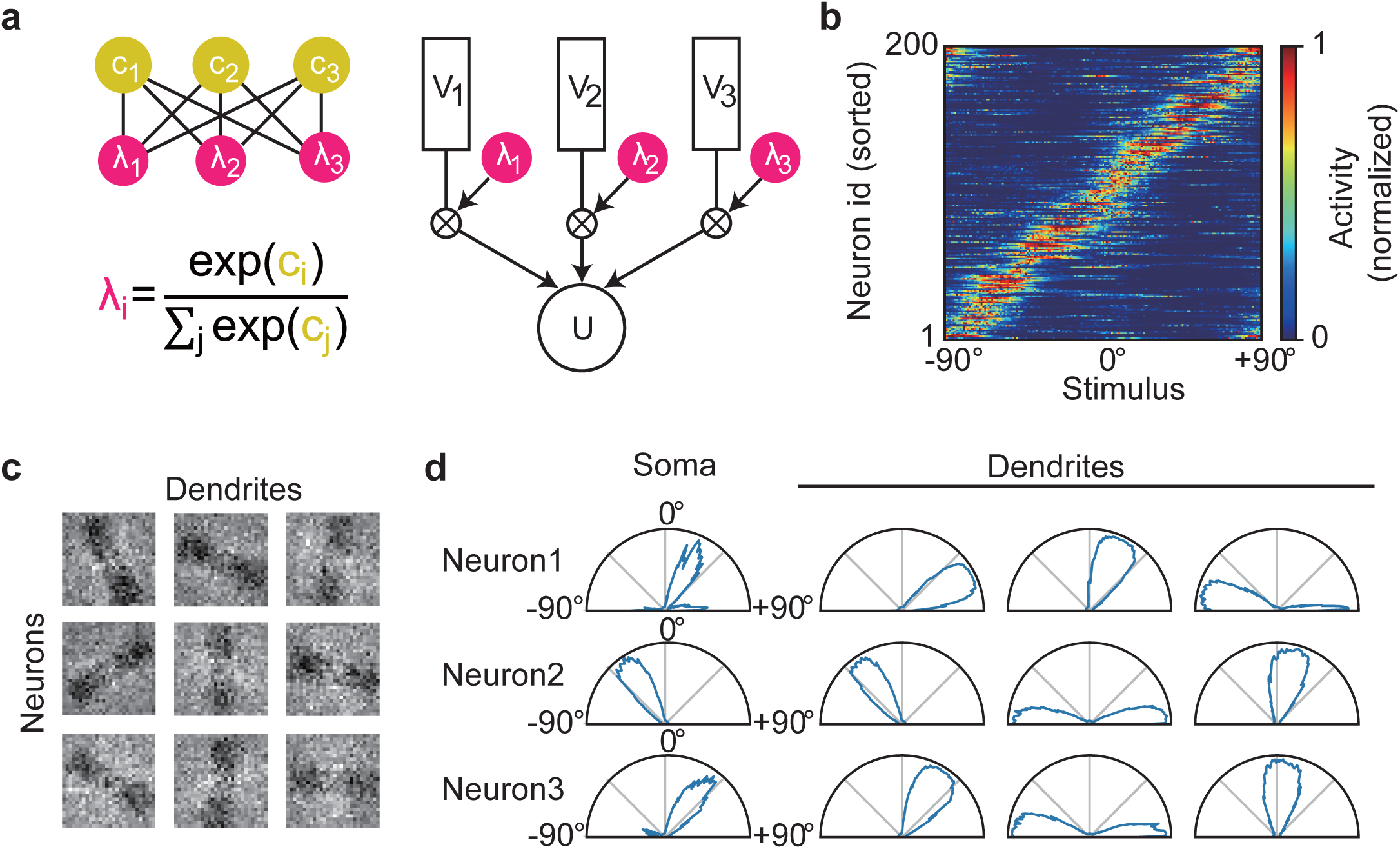
Redundant dendritic representations of preferred sensory features. **a**, A schematic illustration of the neuron model with three dendritic compartments. The dendritic branches have independent gating factors, which compete with each other by softmax. **b**, Somatic responses are shown for all neurons in the trained network. **c**, Trained weight matrices are displayed for afferent inputs to three dendritic branches of three example neurons. **d**, Somatic and dendritic activities of the three neurons in **c** are shown.

### Role of the conventional recurrent synaptic input

While the multiplicative recurrent input (i.e., recurrent gating) is crucial for segregating complex chunks, what is the role of additive (i.e., conventional) recurrent input? We demonstrate that the additive component is still needed for retrieving chunked sequences, namely, for pattern completion. We simulated a network model having both recurrent gating and non-vanishing additive recurrent inputs. The network received a temporal input containing two mutually overlapping chunks (Fig. 8a), and all synaptic connections underwent learning. The trained network formed two cell assemblies responding selectively to either of the chunks, as in the previous network without additive recurrent connections (Supplementary Fig. 5). Since only additive recurrent input, but not recurrent gating, can activate postsynaptic cells, the additive input generates a reverberating activity, which may in turn assist the retrieval of learned sequences. This actually occurred in our simulations. Applying a cue stimulus, which was the first component pattern of one of the learned chunks, enabled the trained network to retrieve the subsequent component patterns in the chunk (Fig. 8b, c). Thus, recurrent gating and additive recurrent inputs contribute to learning and retrieval of segmented sequence memory, respectively, in our network model.

**Figure 8.**
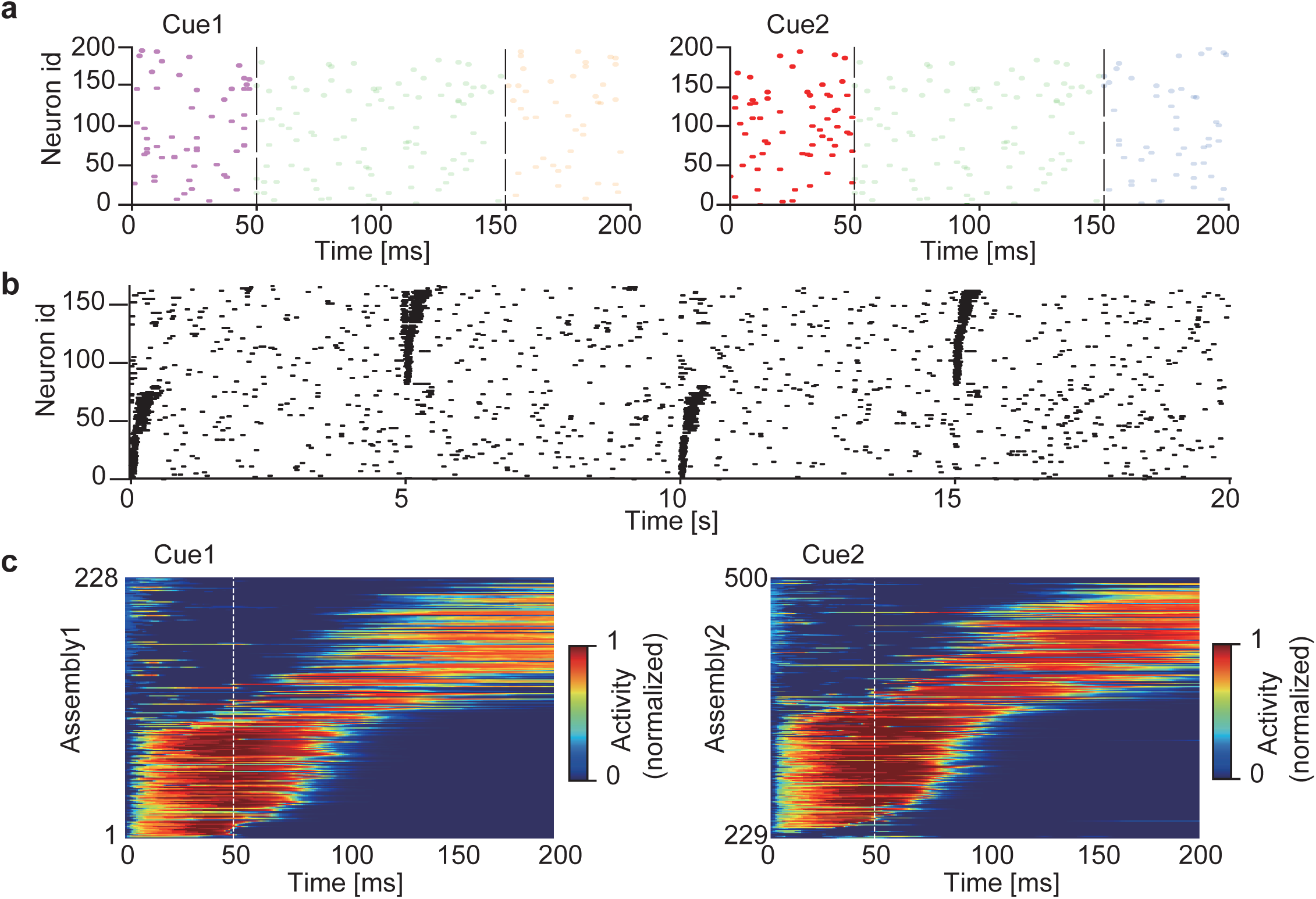
Spontaneous completion of learned sequences. The two chunks shown in Fig. 2 were used for training a network having both recurrent gating and additive recurrent inputs. **a**, In the testing phase, the first component pattern of each chunk (cue 1 or cue2; dark raster plots), but not the subsequent component patterns (light raster plots), was presented to the network. **b**, The raster plot of network activities in the testing phase is shown. Cue 1 and cue 2 were presented alternately every 5 seconds. Neurons were sorted according to their onset response times, and only 160 out of the total 500 neurons are shown for the visualization purpose. **c**, Sequential responses of the two assemblies were averaged over 20 trials. Vertical lines indicate the end of cue presentation. The sequential responses were evoked in the learned order.

## Discussion

In this study, we constructed a recurrent network of compartmentalized neuron models to explore the neural mechanisms to segment temporal input. The crucial role of recurrent gating in context-dependent chunking of complex sequences is a major finding. Recurrent gating enables the instantaneous network state to regulate the degree of the dendro-somatic information transfer in single neurons in different contexts. In contrast, simple segmentation tasks do not necessarily require recurrent connections. With the help of recurrent gating, the model is capable of detecting the fine structures of cell assemblies in large-scale neural recording data.

Our model describes a possible form of integrating dendritic computation into computation at the network level. Learning in our model minimizes the prediction error between the soma and dendrites, thus improving the consistency in responding to synaptic input between the input terminal (dendrites) and the output terminal (soma) of single neurons. This enables the neurons to learn repeated patterns in synaptic input in a self-supervised manner. Previous theoretical studies utilized the local dendritic potential with a fixed gating factor to predict the somatic spike responses [19,21]. We extended the previous learning rule over recurrent connections such that recurrent input helps the dendritic compartment to predict the somatic responses by regulating the degree of signal transfer to the soma in a network state-dependent manner. As a consequence of recurrent gating, the soma can respond differently to the same sequence component depending on the preceding element in sequences, whereas the dendrites respond similarly to the same component. As in our model, some neurons in the premotor nucleus HVC in canaries change their responses to a song element depending on the preceding phrases in songs [13]. However, the response of HVC neurons can also vary according to the following phrases in songs. Such response modulations are likely to represent action planning, which was not considered in this study. Previous experimental and theoretical studies suggested that dendritic inhibition implements a gating operation on synaptic input [29-32]. The role of inhibition on the context-dependent segmentation of input should also be investigated further.

Previously, spike-timing-dependent plasticity was used for detecting recurring patterns in input spike trains in a recurrent neural network without recurrent gating [18]. While the model successfully discriminated relatively simple sequences, it could not discriminate complex sequences involving, for instance, overlapping spatiotemporal patterns. Our results suggest that additive recurrent connections are unlikely to be crucial for learning hierarchically organized sequences. These connections are necessary for retrieving chunked sequences but are unnecessary for learning these sequences. Our results instead suggest that such learning crucially relies on recurrent gating and its context-dependent tunning. Thus, multiplicative and additive recurrent connections have a clear division of labor in the present model. Recurrent synapses were shown to amplify the responses of cortical neurons having similar receptive fields [33], and this amplification resembles the selective amplification of a particular sequence component shown in this study. However, the experimental confirmation of recurrent gating is open to future studies.

Some neural network models in artificial intelligence also utilize gating operations. A well-known example is Long Short-Term Memory (LSTM) for sequence learning and control [34]. The most general form of LSTM contains three types of gate functions, i.e., input, output, and forget gates, and these functions are optimized through supervised learning. In an interesting attempt at the learning-to-learn paradigm [35-37], LSTM was coupled with another network model for self-supervised learning of visual features [38]. In another LSTM-inspired model, neural dynamics with oscillatory gated recurrent input were used to convert spatial activity patterns to temporal sequences in working memory and motor control [22]. In contrast to LSTM, our model learns the optimal gating of the dendro-somatic information transfer by seeking a self-consistent solution to the optimization problem without supervision. Unlike in LSTM, our model only has a single type of gate, which likely corresponds to the input gate of LSTM, to regulate a current flow into the output terminal (soma) of the neuron. In LSTM, however, the most influential gate on learning performance is thought to be the forget gate [39,40]. It is intriguing to ask whether and how recurrent networks learn an optimal forget gate for the unsupervised learning of hierarchical sequences.

When multiple dendritic branches compete for repeated patterns of synaptic input to a single neuron, recurrent gating enables these branches to learn different input features. Consequently, in each neuron, the dendrites learn more redundant representations of input information than the soma. In pyramidal neurons in the rodent primary visual cortex, the dendritic branches have heterogeneous orientation preferences while the somata have unique orientation preferences [26]. Similarly, in retrosplenial cortex [27] and place cells in the hippocampal CA3 [28], the dendritic branches have multiple receptive fields whereas the somata have unique receptive fields. Our model provides a possible neural mechanism for these redundant representations on the dendrites. When the environment suddenly changes, such redundancy may allow neural networks to quickly remodel their responses to adapt to the novel situation. However, the functional benefit of this redundancy has yet to be clarified.

A practically interesting feature of our model is its applicability to large-scale neural recording data. For such purposes, various mathematical tools have been proposed based on methods in computer science and machine learning [41-45]. However, many of these methods suffer time-consuming, combinatorial problems necessary for an exhaustive search for activity patterns in the neural population. In contrast, our model with a biologically inspired learning rule is free from this problem, presumably due to the same reason that cortical circuits do not have this problem. Actually, the present data from the mice visual cortex contain more than 6,000 active neurons, yet our analysis revealed clear evidence for cell assembly structures. These results are interesting because they suggest that cell assemblies underly the multidimensional neural representations of mice spontaneous behavior [24]. As the size of neural recording data is increasing rapidly, the low computational burden and high sensitivity to structured activity patterns show big advantages of this model.

## Methods

### Neural network model

Our network model consists of *N*_in_ input neurons and *N* recurrently connected neurons. Each neuron in the recurrent network consists of two compartments: the somatic and dendritic compartments. Inspired from a previous single neuron model, the somatic response can be approximated as an attenuated version of the dendritic potential *V* [21]. In our recurrent network model, the dendro-somatic signal transfer is regulated by the gating factor *λ* that depends on recurrent synaptic inputs through the local potential *c* as follows:

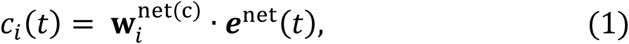

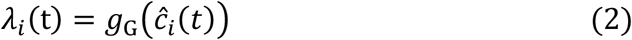

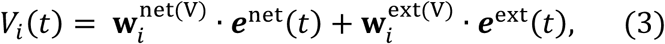

where the subscript *i* is the neuron index, 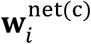 are the *N*-dimensional weight vector of recurrent gating on the local potential *c*, and 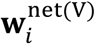 the *N*-dimensional weight vector of additive recurrent connections on the dendrite of the *i*-th neuron. In Eq. (2), *g*_*G*_ and *ĉ* will be defined later. The *N*_in_-dimensional vector 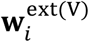 represents the weights of afferent inputs. Except in Fig. 8, we set as 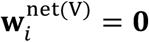. The variables ***e***^net^ and ***e***^ext^ are the post-synaptic potentials evoked by recurrent and afferent inputs, respectively. The initial values of 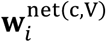 and 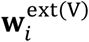 were generated by Gaussian distributions with zero mean and the standard deviations of 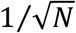 and 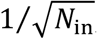, respectively. All three types of connections are fully connected.

The dynamics of the somatic membrane potential are described as

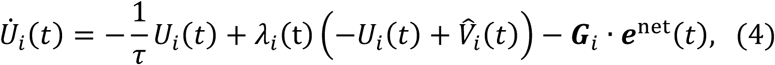

where *τ* = 15 ms, and *ĉ* _*i*_ and 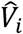 are the standardized potentials calculated as

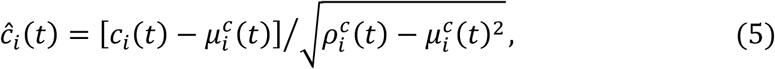

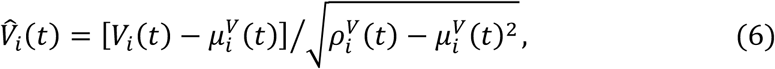

where 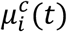 and 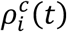 are exponentially decaying averages of the membrane potential and its square of the gating compartment,

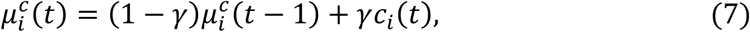

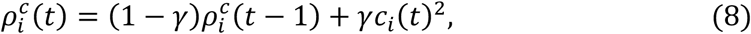

respectively (0 < *γ* < 1). The values of 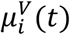 and 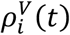 are calculated from *V*_*i*_ in a similar fashion. The last term in Eq. (4) represents peri-somatic recurrent inhibition with uniform inhibitory weights of strength 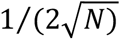. No self-inhibition is considered. In Eq. (2), the gating function *g*_G_(*x*) is defined as

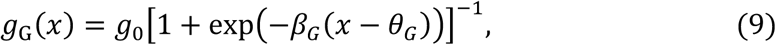

where *g*_0_ = 0.7, *β*_G_ = 5 and *θ*_G_ = 0.5.

The somatic compartment generates a Poisson spike train with saturating instantaneous firing rate given as

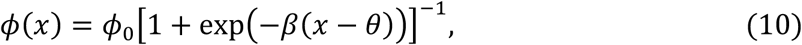

where *ϕ*_0_ = 0.05 kHz, *β* = 5 and *θ* = 1 throughout the present simulations.

Afferent inputs are described as Poisson spike trains of *N*_in_ input neurons:

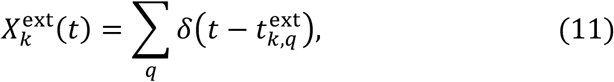

where *δ* is the Dirac’s delta function and 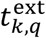 is the time of the *q*-th spike generated by the *k*-th input neuron. The postsynaptic potential evoked by the *k*-th input is calculated as

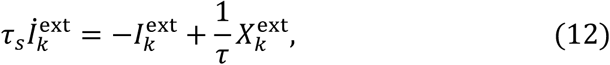

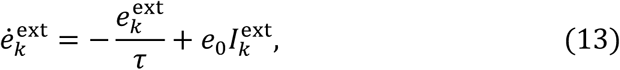

where *τ*_s_ = 5 ms and *e*_0_ = 25. The postsynaptic potentials induced by recurrent inputs, ***e***^net^, are similarly calculated.

### The optimal learning rule for recurrent gated neural networks

We derive an optimal learning rule for the gating recurrent neural network in the spirit of minimization of regularized information loss (MRIL), which we recently proposed for single neurons [19]. The objective function is the KL-divergence between two Poisson distributions associated with the somatic and dendritic activities:

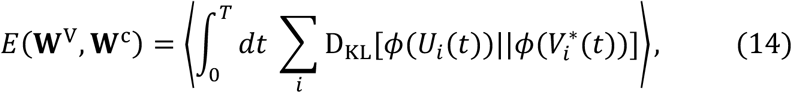

where angle bracket stands for the averaging over input spike trains, and **W**^V^ and **W**^C^ are the weight matrix of synaptic inputs onto the dendrite *V* and those onto the local potential *c* for recurrent gating, respectively. The gated dendritic potential 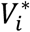 is defined as

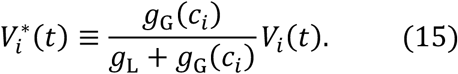

The crucial point in Eq. (15) is that the degree of gating depends on *c*, and hence on network states through Eq. (1).

The learning rule for the weights on the dendritic compartments is similar to the previously derived rule [19] except that the degree of gating is no longer constant in the present model:

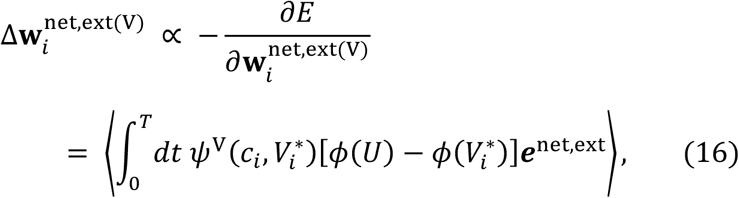

where the function 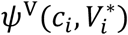 is defined as

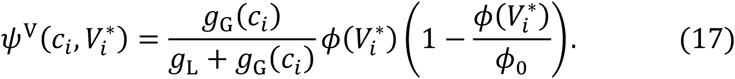

The learning rule for recurrent gating is novel and can be calculated by a gradient descent as follows:

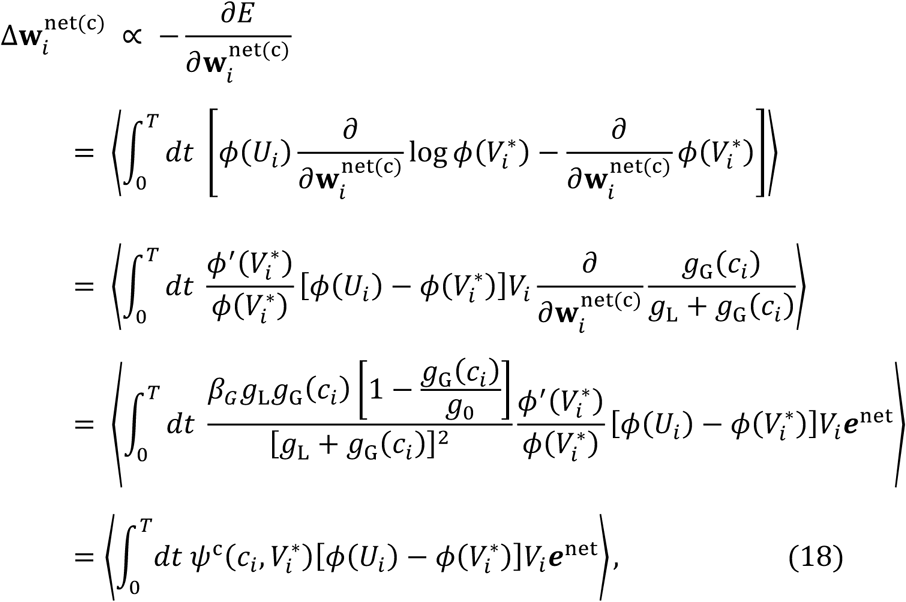

where the function 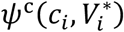 is defined as

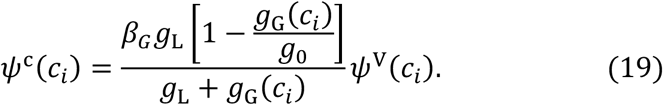

In the present simulations, we used an online version of the above learning rules:

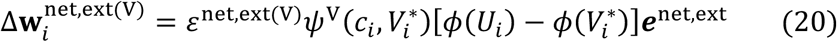

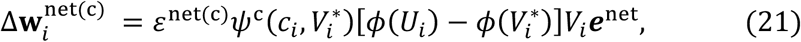

where the learning rates were given as *ε*^ext(V)^ = 10^−5^, *ε*^net(V)^ = 10^−5^, and *ε*^net(c)^ = 10^−4^.

### The optimal learning rule for multi-dendrite neuron model

For the multi-dendrite neuron model used in Fig. 7, the membrane potential of the k-th dendrite of the i-th neuron and the dynamics of the corresponding somatic potential were calculated as

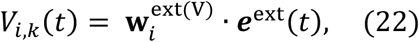

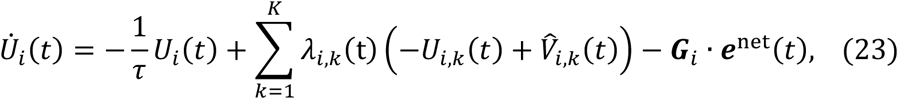

where *K* is the number of dendrites in each neuron. In this study, *K* = 3 for all neurons. We assumed that the dendritic compartments compete for the somatic activity of each neuron, governed by recurrent gating:

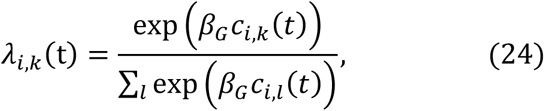

where *c*_*i,k*_ is calculated as

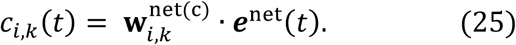

Since 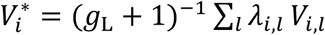, it straightforward to derive the update rule for connections onto to the dendrites:

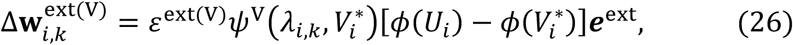

where

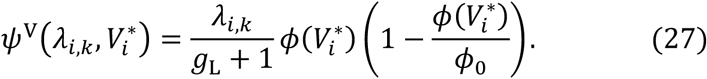

Using the fact that *∂λ*_*i,k*_/*∂c*_*i,l*_ = *λ*_*i,k*_ (*δ*_*l,k*_ − *λ*_*i,k*_), we can derive the update rule for recurrent gating as follows:

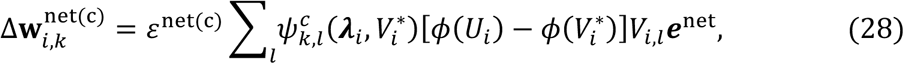

Where

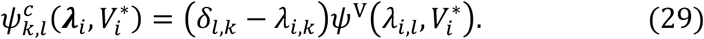

### Normalized mutual information score

In Supplementary Fig. 3, we determined the estimated labels of the output response by Affinity Propagation [46], and then calculated the normalized mutual information score between the estimated labels *X* and the true label *Y* as

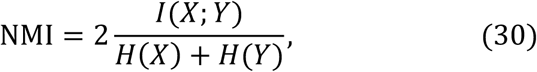

where *I*(*X*; *Y*) is the mutual information between *X* and *Y* and *H*(*X*) is the entropy of *X*.

### The Spearman’s rank-order correlation

In Fig. 5c, we quantified the extent to which the order of sequential responses was preserved in network activity. To this end, we calculated the Spearman’s rank-order correlation between network responses as

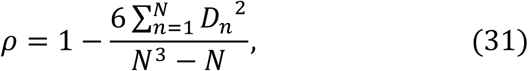

where *N* is the number of neurons in the network and *D*_*n*_ is the difference in the ranks of the n-th neuron between two datasets when sorted according to their onset response times.

### Neural sorting algorithms

In all figures except Fig. 6e, neurons in the trained network were sorted according to the onset response times of these neurons. In Fig. 6e, we first grouped neurons such that all pairs in a group had a correlation coefficient greater than 0.2. We then sorted the resultant groups based on their onset response times. In Fig. 6c and 6f, we first sorted model neurons based on their peak response times. We then sorted the experimental data by associating each cortical neuron with a model neuron showing the highest correlation.

### Values of parameters

The values of parameters used in the present simulations are as follows: in Figs. 2, 4, 5, 8 and Supplementary Figs. 1, 3, and 5, *N* = 500, *N*_in_ = 2000 and *γ* = 0.0003; in Fig. 3 and Supplementary Fig. 2, *N* = 1200, *N*_in_ = 2000 and *γ* = 0.0003; in Fig. 6a-c and Supplementary Fig.4a, *N* = 600, *N*_in_ = 452 and *γ* = 0.0003; in Fig. 6d-f and Supplementary Fig.4b and 4c, *N* = 600, *N*_in_ = 6,532 and *γ* = 0.00005; in Fig. 7, *N* = 200, *N*_in_ = 28 × 28 and *γ* = 0.0003. Usually, the network was trained for the duration of 1,000 seconds. In Fig. 6, the input spike trains constructed from experimental data were repeated 200 times during training.

## Acknowledgements

The authors express their sincere thanks to Thomas Burns for technical assistance. This work was partly supported by KAKENHI (nos. 19H04994) to T.F. T.A was supported by the SRS Research Assistantship of OIST.

## Author contributions

T.F and T.A designed the study and wrote the manuscript, and T.A. performed numerical simulations.

## Competing interests

The authors declare no competing interests.

## Figure legends

**Supplementary Figure 1. Learning of overlapping patterns in a recurrent network without gating. a**, Output spike trains of the trained recurrent network are shown. Neurons were sorted according to their onset response times. **b**, The responses to the two chunks were averaged over 20 trials. **c**, PCA was applied to obtain the low-dimensional trajectories of the trained network. The black, orange, and blue portions indicate the periods of random spike input, chunk 1 and chunk 2, respectively. The two trajectories corresponding to the two chunks were inseparable and the network failed to learn the chunks.

**Supplementary Figure 2. Chunk-selective responses in the network trained in Fig. 3. a**, The responses of the first cell assembly to its preferred (left) and non-preferred (middle, right) chunks are shown. These responses were averaged and normalized as in Fig. 2c. **b**, Preferred responses of two neurons are shown as examples. **c**, As in Fig. 3c-e, trial-averaged responses of dendrite, gating factor and soma are shown for two other neurons.

**Supplementary Figure 3. Robustness against spike timing jitters. a**, Responses of the networks trained on input spike trains with timing jitters of 70 ms (top) and 100 ms (bottom) are shown. Here, spike times within chunks were sifted by the amounts drawn by a Gaussian distribution with mean zero and s.d of jitter strength, and these jitters were present during learning and testing. Neurons were sorted according to the times of their response onsets during chunks, and only 160 out of the total 500 neurons are shown for the visualization purpose. **b**, The normalized average activities of the two assemblies with timing jitters of 70 ms (left) and 100 ms (right)are shown. **c**, Learning curves are shown when the average jitter was 0 ms (purple), 70 ms (green), and 100 ms (blue), respectively. The solid lines and shaded areas represent the averages and s.d over 20 trials, respectively. Learning performance was measured by the normalized mutual information between network activity and target labels (Methods). **d**, The performance measures averaged over 20 trials are shown at various sizes of jitters. Error bars stand for the s.d.

**Supplementary Figure 4. Analysis of calcium imaging data without recurrent gating. a**, The positions of a mouse on a linear track [23] (top) and the activities of model neurons learned without recurrent gating (bottom) are shown. Model neurons were sorted according to their onset response times. Separations between the two sequences corresponding to forward and backward runs are invisible (c.f. Fig. 6b). **b**, Activities of model neurons trained on the neural data recorded from the mice visual cortex [24,25] without recurrent gating are shown. The model neurons were sorted according to their onset response times. **c**, We associated each cortical neuron with a model neuron having the highest correlation with the cortical neuron. Then, we sorted the cortical neurons according to the serial order of model neurons shown in **b**.

**Supplementary Figure 5. Sequence learning in the copresence of recurrent gating and recurrent input**. Dendrites received additive recurrent inputs as well as afferent inputs and the dendritic activity underwent recurrent gating. **a**, As in Fig. 2, the trained network segmented two overlapping chunks in the presence of additive recurrent inputs. **b**, Normalized average responses of two emergent assemblies during the presentations of chunk 1 and chunk 2. **c**, PCA showed that the different chunks were distinguishable by different low-dimensional trajectories, of which the black, orange, and blue portions indicate the periods of random spike input, chunk 1 and chunk 2, respectively.

## Notes

### Competing Interest Statement

The authors have declared no competing interest.

## References

1. Saffran, J.R., Aslin, R.N. & Newport, E.L. Statistical learning by 8-month-old infants. Science 274, 1926–1928 (1996).

2. Wiestler, T. & Diedrichsen, J. Skill learning strengthens cortical representations of motor sequences. Elife 2, e00801 (2013).

3. Waters-Metenier, S., Husain, M., Wiestler, T. & Diedrichsen, J. Bihemispheric transcranial direct current stimulation enhances effector-independent representations of motor synergy and sequence learning. J. Neurosci. 34, 1037–1050 (2014).

4. Dehaene, S., Meyniel, F., Wacongne, C., Wang, L. & Pallier, C. The neural representation of sequences: from transition probabilities to algebraic patterns and linguistic trees. Neuron 88, 2–19 (2015).

5. Leonard, M.K., Bouchard, K.E., Tang, C. & Chang, E.F. Dynamic encoding of speech sequence probability in human temporal cortex. J. Neurosci. 35, 7203–7214 (2015).

6. Naim, M., Katkov, M., Recanatesi, S. & Tsodyks, M. Emergence of hierarchical organization in memory for random material. Sci. Rep. 9, 1–10 (2019).

7. Henin, S. et al. Learning hierarchical sequence representations across human cortex and hippocampus. Sci. Adv. 7, eabc4530 (2021).

8. Franklin, N.T., Norman, K.A., Ranganath, C., Zacks, J.M. & Gershman, S.J. Structured Event Memory: A neuro-symbolic model of event cognition. Psychol. Rev. 127, 327–361 (2020).

9. Miller, G.A. The magical number seven, plus or minus two: Some limits on our capacity for processing information. Psychol. Rev. 101, 343–52 (1956).

10. Ericcson, K.A., Chase, W.G. & Faloon, S. Acquisition of a memory skill. Science 208, 1181–1182 (1980).

11. Orbán, G., Fiser, J., Aslin, R.N. & Lengyel, M. Bayesian learning of visual chunks by human observers. Proc. Natl Acad. Sci. USA 105, 2745–2750 (2008).

12. Christiansen, M.H. & Chater, N. The now-or-never bottleneck: A fundamental constraint on language. Behav. brain sci. 39, e62 (2016).

13. Cohen, Y. et al. Hidden neural states underlie canary song syntax. Nature 582, 539– 544 (2020).

14. Baldauf, D., Cui, H. & Andersen, R.A. The posterior parietal cortex encodes in parallel both goals for double-reach sequences. J. Neurosci. 28, 10081–10089 (2008).

15. Tanji, J., and Shima, K. Role for supplementary motor area cells in planning several movements ahead. Nature 371, 413–416 (1994).

16. Yokoi, A. and Diedrichsen, J. Neural organization of hierarchical motor sequence representations in the human neocortex. Neuron 103, 1178–1190 (2019).

17. Ding, N., Melloni, L., Zhang, H., Tian, X. and Poeppel, D. Cortical tracking of hierarchical linguistic structures in connected speech. Nat. Neurosci. 19, 158–164 (2016).

18. Klampfl, S. & Maass, W. Emergence of dynamic memory traces in cortical microcircuit models through STDP. J. Neurosci. 33, 11515–11529 (2013).

19. Asabuki, T. & Fukai, T. Somatodendritic consistency check for temporal feature segmentation. Nat. Commun. 11, 1554 (2020).

20. Branco, T., Clark, B.A. & Häusser, M. Dendritic discrimination of temporal input sequences in cortical neurons. Science 329, 1671–1675 (2010).

21. Urbanczik, R. & Senn, W. Learning by the Dendritic Prediction of Somatic Spiking. Neuron 81, 521–528 (2014).

22. Heeger, D.J. & Mackey, W.E. Oscillatory recurrent gated neural integrator circuits (ORGaNICs), a unifying theoretical framework for neural dynamics. Proc. Natl Acad. Sci. USA 116, 22783–22794 (2019).

23. Rubin, A. et al. Revealing neural correlates of behavior without behavioral measurements. Nat. Commun. 10, 4745 (2019).

24. Stringer, C. et al. Spontaneous behaviors drive multidimensional, brainwide activity. Science 364, 255 (2019).

25. Stringer, C., Pachitariu, M., Reddy, C., Carandini, M. & Harris, K.D. Recordings of ten thousand neurons in visual cortex during spontaneous behaviors. figshare https://doi.org/10.25378/janelia.6163622.v4 (2018).

26. Jia, H., Rochefort, N.L., Chen, X. & Konnerth, A. Dendritic organization of sensory input to cortical neurons in vivo. Nature 464, 1307–1312 (2010).

27. Voigts, Jakob. & Harnett, M.T. Somatic and dendritic encoding of spatial variables in retrosplenial cortex differs during 2D navigation. Neuron 105, 237–245 (2020).

28. Shannon, K. et al. The dendritic spatial code: branch-specific place tuning and its experience-dependent decoupling. bioRxiv 2020.01.24.916643 (2020).

29. Wang, X.J., Tegner, J., Constantinidis, C. & Goldman-Rakic, P.S. Division of labor among distinct subtypes of inhibitory neurons in a cortical microcircuit of working memory. Proc. Natl Acad. Sci. USA 101, 1368–1373 (2004).

30. Jadi, M., Polsky, A., Schiller, J. & Mel, B.W. Location-dependent effects of inhibition on local spiking in pyramidal neuron dendrites. PLoS Comput. Biol. 8, e1002550 (2012).

31. Sridharan, D. & Knudsen, E.I. Selective disinhibition: a unified neural mechanism for predictive and post hoc attentional selection. Vision Res. 116, 194–209 (2015).

32. Yang, G.R., Murray, J.D. & Wang, X.J. A dendritic disinhibitory circuit mechanism for pathway-specific gating. Nat. Commun. 7, 12815 (2016).

33. Peron, S. et al. Recurrent interactions in local cortical circuits. Nature 579, 256–259 (2020).

34. Hochreiter, S. & Schmidhuber, J. Long short-term memory. Neural Comput. 9, 1735–1780 (1997).

35. Schmidhuber, J., Zhao, J. & Wiering, M. Shifting inductive bias with success-story algorithm, adaptive Levin search, and incremental self-improvement. Machine Learning. 28, 105–130 (1997).

36. Thrun, S. & Pratt, L. Learning to learn: Introduction and overview. Learning to learn. (Springer, Boston, MA, 1998).

37. Finn, C., Abbeel, P. & Levine, S. Model-agnostic meta-learning for fast adaptation of deep networks. In Int. Conf. Machine Learning 1126-1135 (ML Research Press, 2017).

38. Wortsman, M., Ehsani, K., Rastegari, M., Farhadi, A. & Mottaghi, R. Learning to learn how to learn: Self-adaptive visual navigation using meta-learning. In Proc. IEEE/CVF Conference on Computer Vision and Pattern Recognition. 6750-6759 (IEEE, 2019).

39. Gers, F.A., Schmidhuber, J. & Cummins, F. Learning to Forget: Continual Prediction with LSTM. Neural Comput. 12, 2451–2471 (2000).

40. Greff, K., Srivastava, R.K., Koutník, J., Steunebrink, B. R. & Schmidhuber, J. LSTM: A search space odyssey. IEEE Trans. Neural Netw. 28, 2222–2232 (2016).

41. Abeles, M., Bergman, H., Margalit, E. & Vaadia, E. Spatiotemporal firing patterns in the frontal cortex of behaving monkeys. J. Neurophysiol. 70, 1629–1638 (1993).

42. Euston, D.R., Tatsuno, M. & McNaughton, B.L. Fast-forward playback of recent memory sequences in prefrontal cortex during sleep. Science 318, 1147–1150 (2007).

43. Shimazaki, H, Amari, S., Brown E.N. & Grün, S. State-space analysis of time-varying higher-order spike correlation for multiple neural spike train data. PLoS Comput. Biol. 8, e1002385 (2012).

44. Mackevicius, E.L. et al. Unsupervised discovery of temporal sequences in high-dimensional datasets, with applications to neuroscience. Elife 8, e38471 (2019).

45. Watanabe, K., Haga, T., Tatsuno, M., Euston, D.R. & Fukai, T. Unsupervised detection of cell-assembly sequences by similarity-based clustering. Front. Neuroinform 13, 39 (2019).

46. Frey, B.J. & Dueck, D. Clustering by passing messages between data points. Science 315, 972–976 (2007).

