## Supplementary figures and images for "Neural mechanisms of context-dependent segmentation tested on large-scale recording data"

### Supplementary Figure 1

#### Supplementary Figure 1

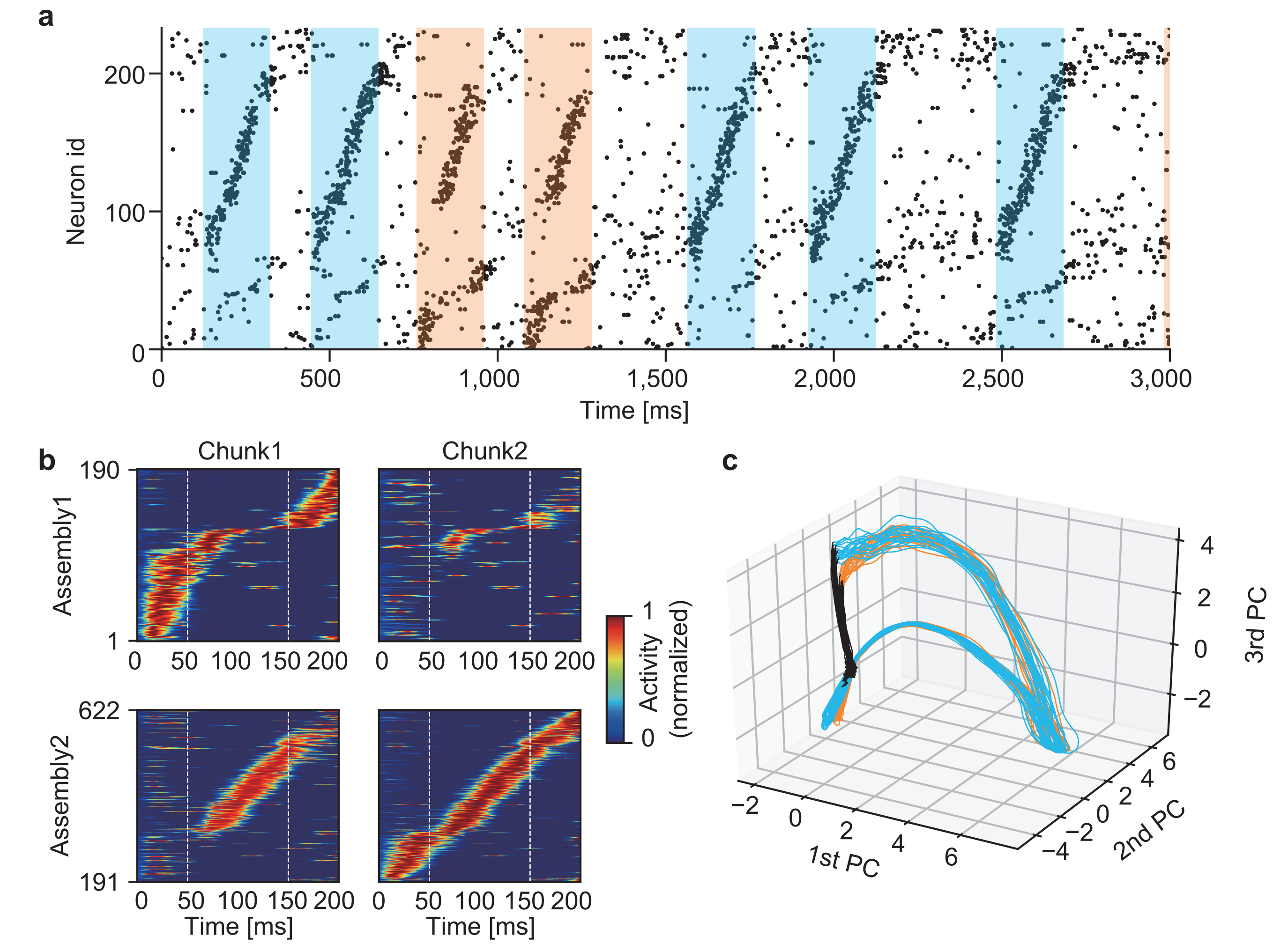

### Supplementary Figure 2

**Supplementary Figure 2**

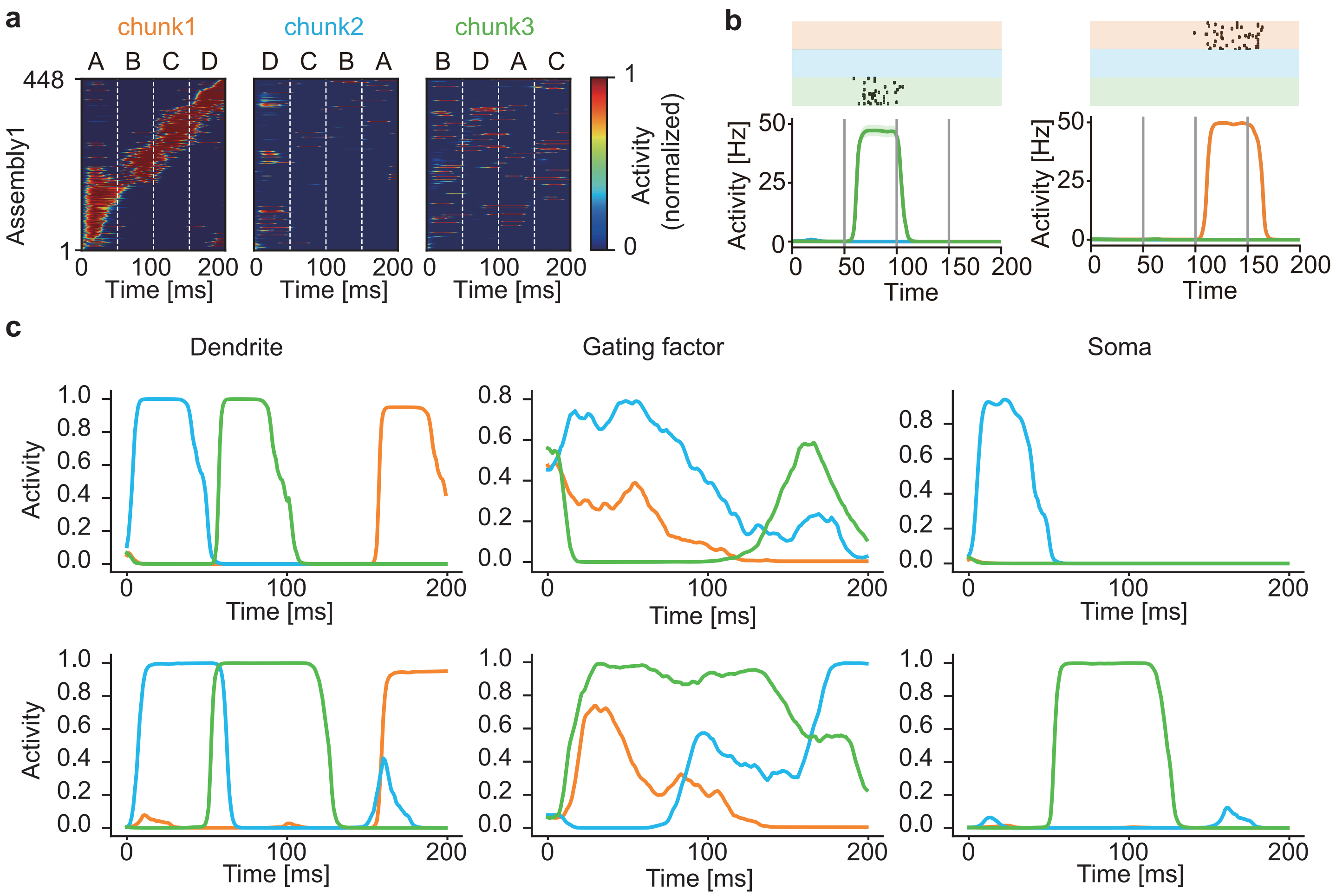

### Supplementary Figure 3

**Supplementary Figure 3**

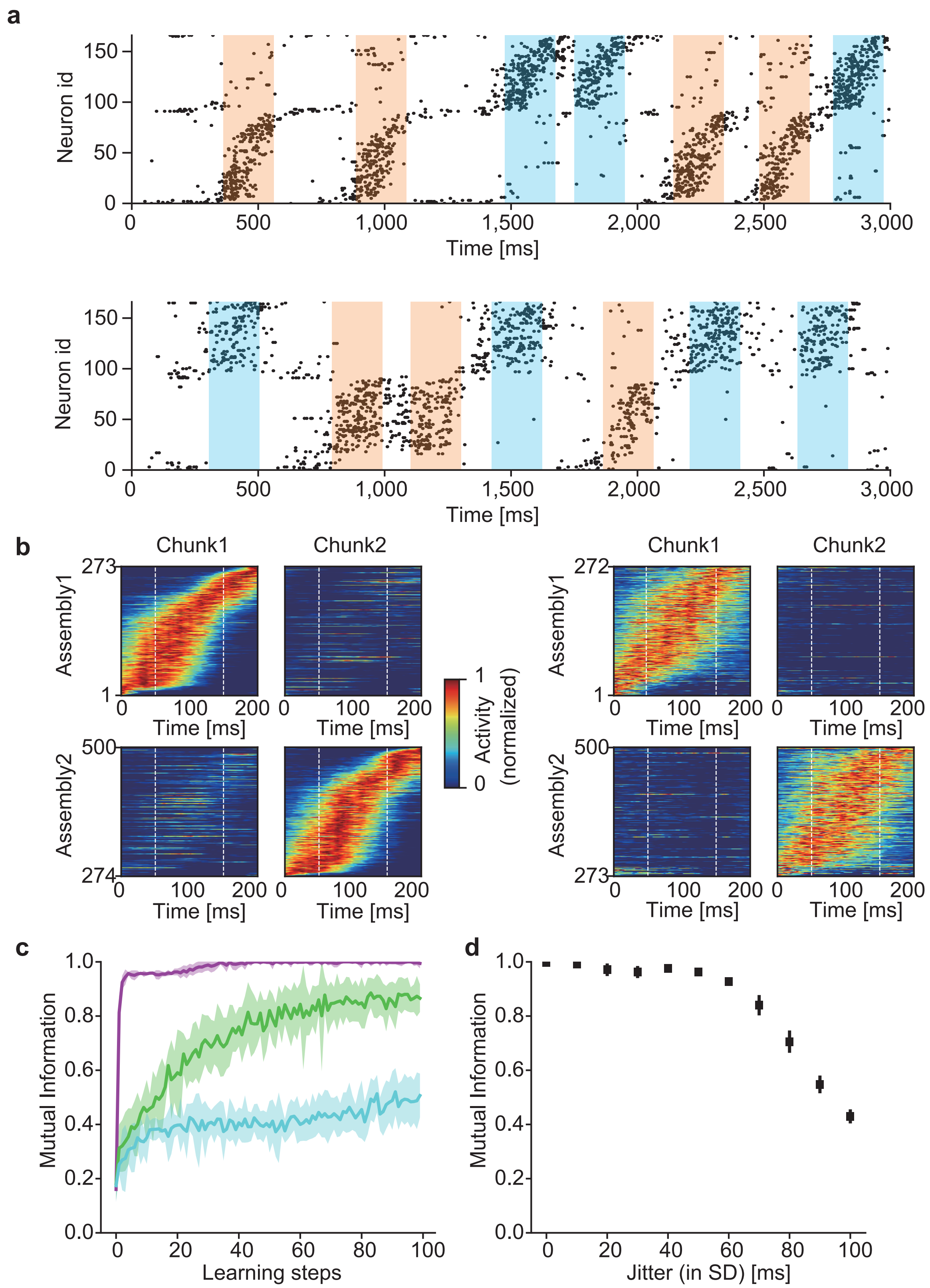

### Supplementary Figure 4

#### Supplementary Figure 4

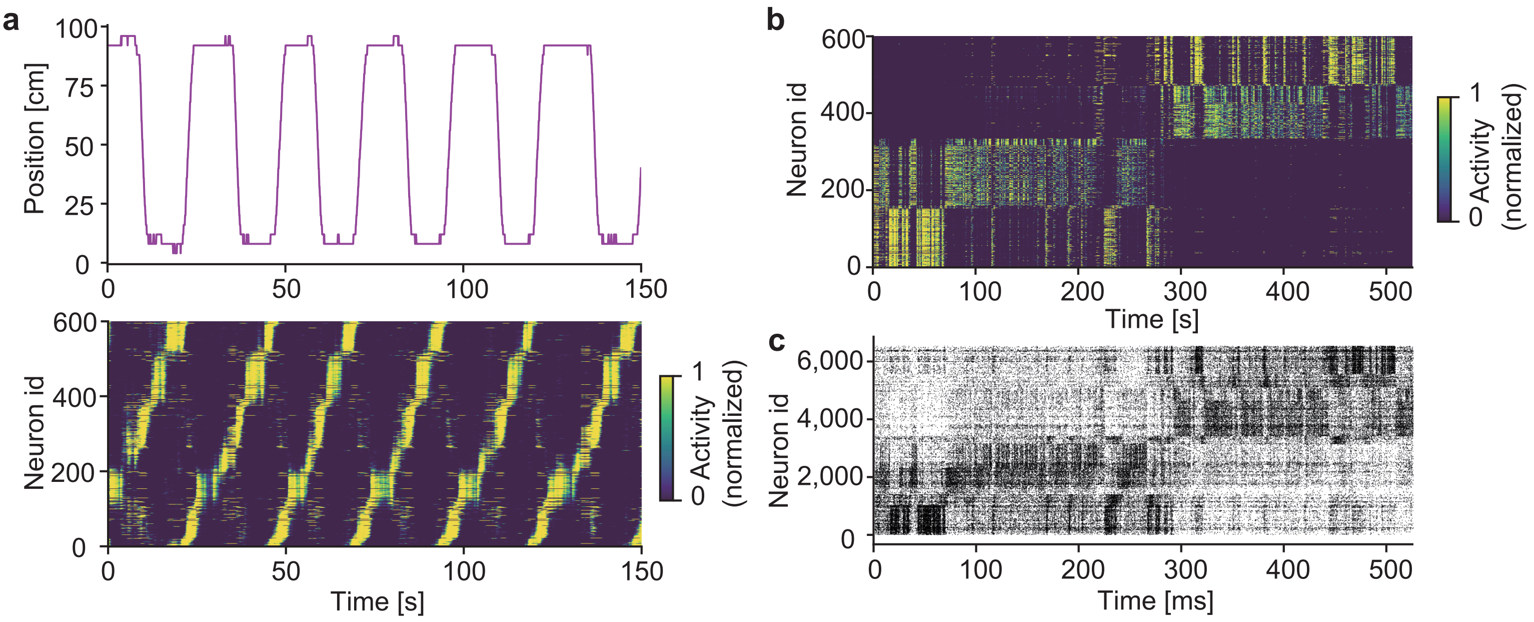

### Supplementary Figure 5

**Supplementary Figure 5**

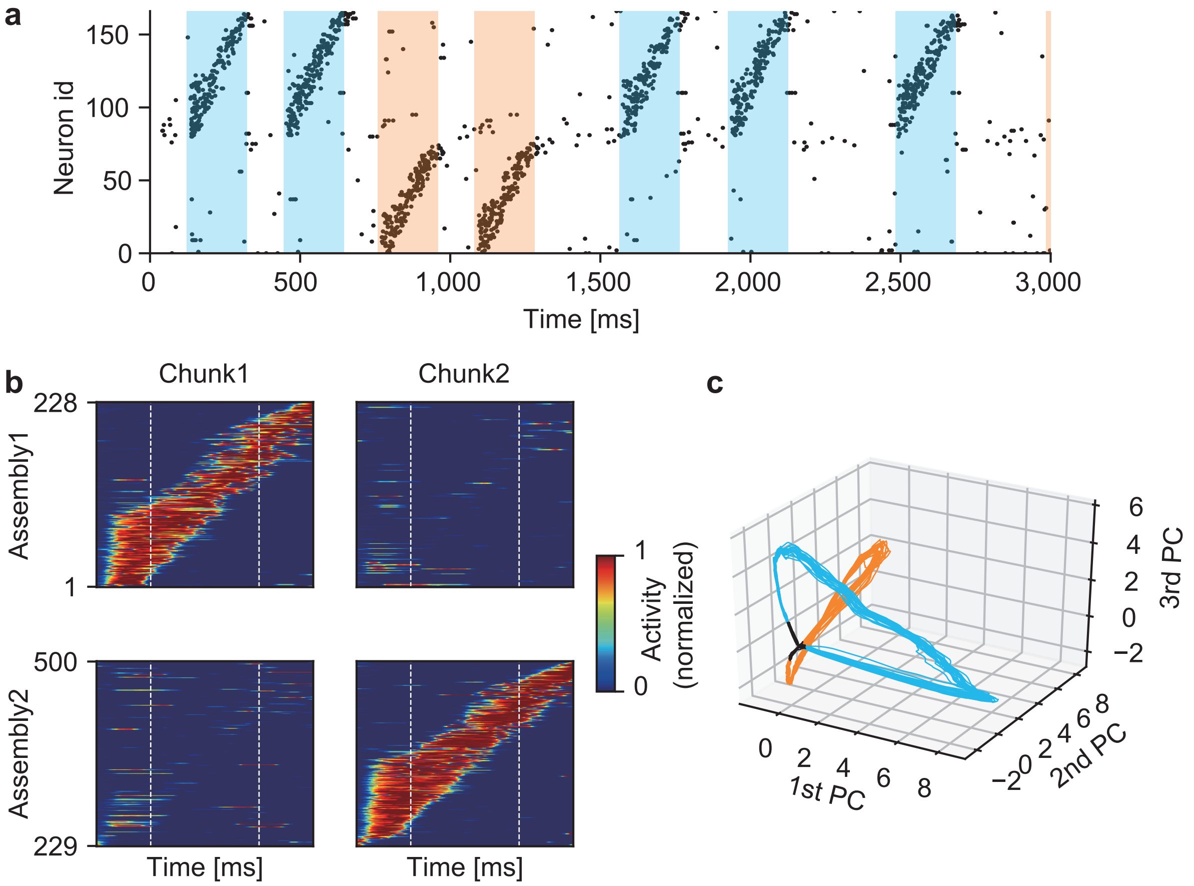
